# The Endogenous Opioid Met-Enkephalin Modulates Thalamo-Cortical Excitation Inhibition Balance in a Medial Thalamus-Anterior Cingulate Cortex Circuit

**DOI:** 10.1101/2023.07.13.547220

**Authors:** Erwin Arias Hervert, William Birdsong

## Abstract

Activation of opioid receptors in the anterior cingulate cortex (ACC) mediates aspects of analgesia induced by both exogenous and endogenous opioids. We have previously shown that opioid signaling disrupts both afferent excitatory and indirect inhibitory synaptic transmission from the medial thalamus (MThal) to the ACC, but the effects of endogenous opioids within this circuit remain poorly understood. The goal of the current study was to understand how the endogenous opioid, [Met]5-enkephalin (ME), modulates thalamic-driven excitatory and inhibitory synaptic transmission onto layer V pyramidal neurons in the ACC. We used pharmacology, brain slice electrophysiology and optogenetic stimulation to study opioid-mediated modulation of optically evoked glutamatergic and GABAergic transmission. The results revealed that ME inhibited both AMPA-mediated excitatory and GABA-mediated inhibitory synaptic transmission in the ACC. However, inhibitory transmission was more potently inhibited than excitatory transmission by ME. This preferential reduction in GABAA-mediated synaptic transmission was primarily due to the activation of delta opioid receptors by ME and resulted in a net disinhibition of MThal-ACC excitatory pathway. These results suggest that moderate concentrations of ME can lead to net excitation of ACC circuitry and that analgesia may be associated with disinhibition rather than inhibition of ACC subcircuits.

## INTRODUCTION

According to the International Association for the Study of Pain (IASP), pain is an unpleasant sensory and emotional experience that is associated with, or similar to, actual or potential tissue damage (Raja et al. 2020). While acute pain perception is considered necessary for the survival of an organism (Iadarola and Caudle 1997), prolonged pain, which can result from inflammation (Matisz and Gruber 2022), nerve injuries (Alles and Smith 2018) or internal organ damage (Grundy, Erickson, and Brierley 2019), can be detrimental. Noxious stimuli activate cortical and sub-cortical structures such as the anterior cingulate cortex (ACC) and the thalamus (Coghill et al. 1994). Hyperactivity of the ACC has been correlated with acute (Rainville et al. 1997) and chronic pain (Hsieh et al. 1995; J. P. Johansen, Fields, and Manning 2001; Joshua P. Johansen and Fields 2004), whereas manipulations that decreased ACC activity such as cingulotomy (Folt and White 1963; Allam et al. 2022) or optogenetic inactivation (Elina et al. 2021) reduced the unpleasantness of noxious stimuli without modifying nociception.

The medial thalamus (MThal), particularly the mediodorsal nucleus of the thalamus (MD), is a major source of nociceptive information to the ACC. Activation of nociceptive MD neurons by mechanical or thermal noxious stimuli is highly correlated with activation of neurons in the ACC (Lee et al. 2007), whereas inactivation of nociceptive thalamic neurons blocks ACC nociceptive activity (Sikes and Vogt 1992). Furthermore, optogenetic activation of MD presynaptic terminals to the ACC aggravates aversive behaviors in animal models of neuropathic pain (Meda et al. 2019). Thus, MThal-ACC thalamocortical circuits play a central role in modulating pain perception and motivated behaviors in humans and rodents, and it constitutes a potential target for therapeutics to manage the unpleasantness of pain.

Current therapies for management of severe pain rely heavily on prescription opioid drugs, which modulate pain perception with high efficacy (Tobin et al. 2022) through activation of mu opioid receptors (MOR) expressed in neuronal structures involved in pain perception across the nervous system (Corder et al. 2018). Injection of morphine into the ACC has been shown to diminish the aversiveness of pain perception (Navratilova et al. 2015). Delta (DOR) and kappa (KOR) opioid receptors are also expressed in many brain regions involved in pain processing and DOR and KOR agonists also modify pain processing. DOR signaling in the ACC has also been demonstrated to diminish pain-related aversion (Ma et al. 2022). Opioid receptors are G protein-coupled receptors (GPCRs) that can be activated by endogenous or exogenous opioids like [Met]^5^-enkephalin (ME) or morphine, respectively (Paul, Sribhashyam, and Majumdar 2023). Critically, non-opioid analgesics may mediate some effects through opioid receptor activation by endogenous opioids. For example, injection of an opioid receptor antagonist into the ACC attenuates pain relief induced by intrathecally administered analgesics suggesting that endogenous opioids mediate this effect (Navratilova et al. 2015). Understanding how both endogenous and exogenous opioids modulate specific neuronal circuits involved in pain perception is essential to designing future analgesic drugs without the adverse effects of typical opioids such as analgesic tolerance and hyperalgesia (Mercadante, Arcuri, and Santoni 2019; Colvin, Bull, and Hales 2019), constipation and bowel dysfunction (Farmer et al. 2018), respiratory depression and death (Bateman, Saunders, and Levitt 2023).

MThal axons form axo-dendritic synaptic connections with pyramidal neurons located in layers II/III, V, and VI of ACC (Georgescu, Popa, and Zagrean 2020). MThal axons also synapse onto local parvalbumin interneurons which exert powerful di-synaptic feedforward inhibition on pyramidal neurons within the ACC and allow for precise temporal integration of excitatory inputs in the pyramidal cells (Delevich et al. 2015). We have previously shown that activation of MOR potently inhibits both excitatory and feedforward-inhibitory responses elicited by stimulation of MThal synapses in the ACC. In contrast, DOR activation selectively inhibits feedforward inhibitory transmission (Birdsong et al. 2019). Endogenous enkephalins are non-selective opioid agonists that are expressed in the ACC. Because enkephalins can activate both DOR and MOR, they may have effects on both excitatory and inhibitory signaling within the ACC. These effects may depend on the relative sensitivity of MOR and DOR to enkephalin (Mansour et al. 1995; Emery and Akil 2020; Corder et al. 2018) and the relative expression levels of MOR and DOR in both inhibitory and excitatory circuits. In the present study we determined the effect of various concentrations of enkephalin on both excitatory and inhibitory ACC signaling in response to activation of MThal terminals. We found a concentration-dependent effect on modulation; moderate concentrations of enkephalin primarily inhibited inhibitory signaling, leading to ACC disinhibition, while higher concentrations of enkephalin inhibited both excitation and inhibition. Thus, enkephalin had a concentration dependent biphasic effect on MThal-ACC circuitry in a manner that depended on the relative activation of DOR and MOR within the ACC.

## METHODS

### Ethical approval

All animal handling and experimental procedures associated with or performed in this study followed National Institutes of Health (NIH) animal use guidelines and were approved by the Institutional Animal Care & Use Committee (IACUC) at the University of Michigan (Approval Number PRO00010677). All investigators understand the ethical principles under which Neuropharmacology works, and the work follows the journal’s animal ethics checklist, Animals were housed in an enriched environment in a 12 hr light–12 hr dark cycle and climate-controlled room (22°C) with free access to water and food. All research performed in this study followed the Institutional Biosafety Committee at the University of Michigan (Approval Number IBCA00001255).

### Animals

Both male and female mice between 6-12 weeks old at the time of the electrophysiological recordings were used for the experiments presented in this study. C57/BL6J mice were obtained from The Jackson Laboratory (JAX) and bred in our laboratory. Breeding, husbandry and genotyping was performed by designated personnel in the animal care facility of the Pharmacology Department at the University of Michigan.

### Stereotactic microinjection of viral vectors

Mice (4-10 weeks old) were placed in an induction chamber and anesthetized with 5% isoflurane by inhalation, then transferred to the stereotactic apparatus where 2% isoflurane was delivered through a nosecone for maintenance and 5 mg/kg carprofen was injected subcutaneously to induce analgesia. The mouse’s head was fixed and stabilized in the stereotactic apparatus (Kopf Instruments, model 1400). The scalp was shaved and sterilized, and a centimeter-long surgical window was made by cutting the scalp with a scalpel along the anteroposterior axis to expose the skull. A craniotomy was performed over the target site bilaterally. AAV2-syn-hChR2(H134R)-EYFP (UNC vector core) was injected at a volume of 64.4 nl in the medial thalamus (MThal) centered around the mediodorsal thalamus (coordinates in mm: AP: -1.1, ML: +/-0.55, DV: -3.6) using a Nanoject II microinjector (Drummond Inc). Coordinates were selected with the aid of an online mouse brain atlas and based on our prior work (Paxinos, 2001; Hunnicutt et al., 2016; Birdsong et al., 2019). Mice were observed for 7 days post-surgery and detailed post-operative surgery records were kept in the laboratory. All surgical tools were sterilized prior to performing the procedure.

### Brain slice preparation for electrophysiology

Three weeks after injection, mice were deeply anesthetized with isoflurane using the drop-jar method and euthanized by cervical dislocation and rapid decapitation. The ventilation rate and pedal reflex were used as indicators of an adequate level of anesthesia before euthanasia. The brain was carefully removed from the skull after decapitation and transferred to a petri dish filled with ice-cold Krebs-Ringer solution (in mM: 136 NaCl, 2.5 KCl, 1.2 MgCl_2_, 2.4 CaCl_2_, 1.2 NH_2_PO_4_, 21.4 NaHCO3, 11.1 Dextrose; 310-320 mOsm/kg) supplemented with 5 mM kynurenic acid and saturated with 5% CO_2_:95% O_2_ gas mixture. 300 μm coronal sections were prepared using a vibratome (7000smz-2, Campden Instruments). Sections were collected and transferred to a recovery chamber containing Krebs solution at room temperature.

Fluorescence and transmitted light images were obtained from live brain slices at both the thalamic injection site and the ACC (Nikon AZ-100 fluorescent microscope) to verify proper injection site placement and expression.

### Patch-clamp electrophysiology

Recordings in voltage-clamp configuration were made using a low chloride cesium-based internal solution (in mM: 135 Cs-gluconate, 1 EGTA, 1.5 MgCl_2_, 10 HEPES, 3 NaCl, 0.4 GTP, 1.8 ATP and 8 mM phosphocreatine; pH=7.4; 290 mOsm/kg). Recordings in current-clamp configuration were performed using a low chloride potassium-based internal solution (in mM: 130 K-gluconate, 5 NaCl, 1.5 MgCl_2_, 10 HEPES, 0.1 EGTA, 0.4 GTP, 1.8 ATP and 8 mM phosphocreatine, 3 QX314; pH=7.4, 280 mOsm/kg). Oxygenated Krebs solution was used as the extracellular medium in both cases. For voltage-clamp experiments, the Krebs solution was supplemented with 3 mM MPEP, 10 mM CGP55845, 30 mM mecamylamine, and 10 mM scopolamine. Fire polished borosilicate pipettes with 3-4 MOhm resistance (Sutter Instruments, Novato, CA) were filled with either Cs-gluconate or K-gluconate internal solutions. Recordings were performed on putative ACC layer V pyramidal neurons characterized visually by their anatomic location, size, and morphology. A 0.5-1 ms duration optical stimulus was delivered using a digitally-controlled LED driver and a 470 nm wavelength LED (Thorlabs, Newton, NJ) through a 60x, 1 NA water immersion objective (Olympus, Tokyo, Japan, BX51W). at both the injection sites and the ACC The stimulation amplitude was adjusted to yield synaptic currents with EPSC amplitudes >0.05nA and IPSC amplitudes <2.0 nA. Approximately 1mW of power exiting the objective was generally necessary to achieve currents meeting these criteria. The series resistance was monitored throughout the experiments and only cells where the series resistance was <20 MOhm were considered for the analysis. The cells where the series resistance changed more than 10% with respect to baseline over the course of the experiment were excluded from the analysis. A stable baseline was obtained in whole-cell configuration by stimulating with single LED pulses at 0.5 Hz for 10 minutes. For voltage clamp recordings, EPSCs and IPSCs were electrically isolated electrically by holding the membrane potential at the IPSC or EPSC reversal potentials, -65 and +5 mV not corrected for junction potential, respectively. The reversal potentials of EPSCs and IPSC with these solutions were found empirically as described previously (Birdsong, 2019).

During recording, slices were continuously perfused with warmed (32-34°C) oxygenated Krebs solution at a rate of 2-3 mL/min. Drug was applied by bath perfusion for 10-15 minutes depending on the compound and dose used followed by a 10–15-minute washout with regular Krebs solution in the experiments with enkephalin or Krebs solution supplemented with 1 μM naloxone for all other agonists. Each slice was used for only a single recording where drug was applied.

### Data acquisition and quantification

Data were acquired by using a Multiclamp 700B amplifier (Molecular Devices) and Wavesurfer (Janelia Research Campus) or Axograph (John Clements) and digitized at 10 kHz with either InstruTECH LIH 8+8 data acquisition system (HEKA) or BNC-2009A/ PCIe-6353 (National Instruments). Data analysis was performed offline using Axograph (John Clements), Google Sheets (Google), and Graphpad Prism (Dotmatics). The average maximum current amplitude was calculated by substracting the baseline of individual sweeps and averaging the last 5 sweeps for baseline, drug, and washout conditions, respectively. The amplitude was calculated as the maximum positive or negative current with respect to baseline for IPSC and EPSC, respectively. The amplitudes were normalized with respect to baseline by dividing the average maximum amplitude in presence of the drug by the average maximum amplitude during baseline. The excitation-inhibition ratio (E-I ratio) was calculated by dividing the average maximum amplitude of the EPSC by the corresponding average maximum amplitude of the IPSC in each condition.

## Statistical analysis

An ordinary two-way ANOVA with multiple comparisons was used to determine the significance of the differences between EPSC and IPSC sensitivities to enkephalin across the range of concentrations tested. EC_50_ values were estimated by non-linear regression fitting the following model to our data:

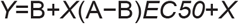

The *EC50* values represent the concentration of agonist that gives a response halfway between minimum and maximum values. *A* and *B* are the maximum and minimum values and represent plateaus in the percentage of inhibition by enkephalin. We constraint the model to *EC50*>0 and *B*=0 (Graphpad Prism Curve Fitting Guide). The differences in E/I balance were determined with a one-way ANOVE with Dunnett’s multiple comparisons test when the data was presented as the Log(ΔE/Ibalance) change from baseline. Multiple paired t tests with individual variance for each group were used to assess statistical significance when data was analyzed as baseline versus drug. Average values were presented as mean ± standard deviation unless otherwise stated, N equals number of animals, while n equals number of cells.

## RESULTS

### Enkephalin modulated thalamic-mediated EPSCs and IPSCs in ACC layer V pyramidal neurons in a concentration-dependent manner

To determine how enkephalin affected thalamo-cortical excitatory and feedforward inhibitory circuits in the ACC, whole-cell patch clamp recordings were obtained from layer V pyramidal neurons in the ACC in acute brain slices prepared from mice expressing channelrhodopsin 2 (ChR2) in MThal projection neurons (Figure 1A). Optically-evoked, electrically-isolated excitatory postsynaptic currents (EPSCs) and inhibitory postsynaptic currents (IPSCs) were recorded in response to excitation of ChR2-expressing MThal terminals in ACC, before and after application of various concentrations of ME (Figure 1B). ME was applied at a range of concentrations from 0.01-3 µM and EPSC and IPSC amplitudes were measured and compared to baseline measurements. ME decreased the peak amplitudes of both EPSCs and IPSCs in a concentration-dependent manner (Figure 1C). The maximum inhibition and the concentration of enkephalin needed to achieve a half maximal effect (EC_50_) were determined for both EPSCs and IPSCs through a non-linear regression analysis. The average maximum inhibition of the EPSC was 48.53% (40.32 to 58.37 percent, 95% CI) while the average maximum inhibition of the IPSC was 72.92% (65.02 to 83.26 percent, 95%CI). The EC50 values were individually calculated to be 0.0894 µM (0.0395 to 0.1981 micromolar, 95% CI) for EPSC and 0.0241 µM (0.0152 to 0.0511, 95% CI) for IPSC. A statistical analysis using a repeated measures (EPSC and IPSC) two-way ANOVA with multiple comparisons showed significant main effects of ME concentration (p<0.0001) and EPSC vs IPSC (p<0.0001) and an interaction between the ME concentration and inhibition of EPSCs vs. IPSCs (p=0.0001). The inhibition of EPSCs and IPSCs by enkephalin was statistically different at concentrations of 0.03, 0.1, 0.3, 1 and 3 µM. The average inhibition for EPSC and IPSC were 20.94±9.29% and 64.72±15.89% of the initial responses at 0.1 µM, respectively (n=9, N=9; ****p<0.0001); 42.14±24.16% and 66.41±25.39% at 0.3 µM (n=9, N=9; *p<0.05); and 40.05±13.96% and 70.73±16.32% at 1 µM (n=8, N=8; **p<0.01, Figure 1D).

**Figure 1:**
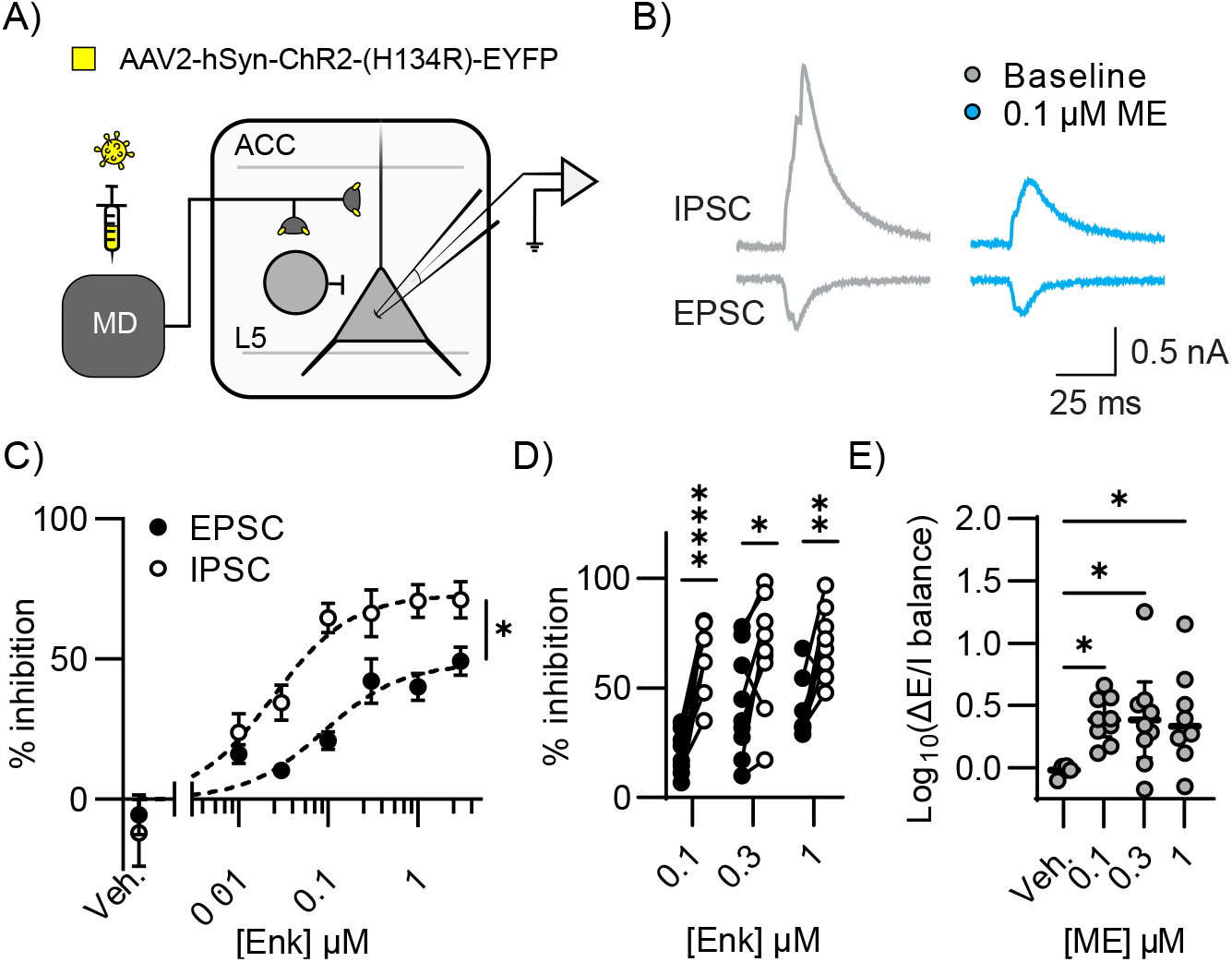
[Met]5-Enekphalin preferentially inhibited polysynaptic MThal-ACC inhibitory signaling. A) Schematic of viral-mediated ChR2 expression in MThal and whole cell voltage clamp recording from layer 5 (L5) ACC pyramidal cells in acute brain slices. B) Representative average optically evoked EPSCs (lower) and IPSCs (upper) recorded under baseline conditions (gray) and in the presence of 0.1 µM ME (blue). C) Concentration response curve generated from EPSC (solid circles) and IPSC (open circles) inhibition as in “B” by various concentrations of ME or vehicle. Data are plotted as % inhibition of the peak EPSC or IPSC relative to baseline. Data were fit with a nonlinear fit in Prism. Probability that the two data sets could be fit with the same parameters: p<0.0001. D) Relative inhibition of pairs of EPSCs (solid) and IPSCs (open) by 0.1, 0.3 and 1 µM ME demonstrating that IPSCs were reliably inhibited more than EPSCs. (two-way ANOVA with Šídák’s multiple comparisons test, ****p<0.0001, **p<0.01, *p<0.05). E) change in excitation/ inhibition (E/I balance) caused by vehicle, 0.1, 0.3, and 1 µM ME calculated as (EPSC/IPSC) _drug_ /(EPSC/IPSC)_baseline_. At 0.1, 0.3 and 1 µM ME the E/I balance increased significantly compared to vehicle (one-way ANOVA with Dunnett’s multiple comparisons, *p<0.05).

The ratio of the EPSC amplitude relative to IPSC amplitude (E/I) ratio can give a rough estimate of net excitatory drive. Because IPSCs were more potently inhibited by ME than EPSCs, we expected that the E/I ratio would increase in the presence of ME. Our results also showed that the synaptic E/I ratio increased significantly when enkephalin was present at concentrations of 0.1, 0.3, and 1 µM (as determined by ordinary one-way ANOVA and Dunnett’s multiple comparisons test). In the presence of vehicle (bestatin/ thiorphan) the mean E/I ratio relative to the baseline E/I ratio did not change (expressed as Log_10_(ΔE/I balance), (Log_10_(ΔE/I balance)=-0.02013±0.01 (n=6, N=2). However, in the presence of ME (0.1-1µM), the mean E/I ratio approximately doubled (Figure 1E) and was significantly greater than the effect of vehicle alone (Log_10_(ΔE/I balance)=0.3865±0.06 (0.1µM ME, n=9, N=9, *p<0.05), 0.3846±0.13 (0.3µm ME, n=9, N=9, *p<0.05) and 0.4054±0.13 (1µM ME, n=9 and n=8, *p<0.05). Taken together, these results demonstrate that ME inhibited both MThal-evoked EPSCs and IPSCs and that IPSCs were more potently inhibited. At 0.1 µM, ME reliably inhibited IPSC amplitude while having a relatively modest effect on EPSC amplitude.

### Enkephalin-induced inhibition of thalamic driven synaptic transmission in the ACC was mediated by both MOR and DOR

As a non-selective opioid agonist, ME can activate both MOR and DOR. To determine the contribution of MOR and DOR to the effects of ME on the EPSC, IPSC and E/I balance, ME was applied to slices in the presence of selective MOR or DOR antagonists. The contribution of MOR activation to the effect of enkephalin was studied by recording the thalamic driven EPSCs and IPSCs in layer V pyramidal neurons in the ACC in the presence of the DOR antagonist TIPP[psi] (1 µM) and ME (0.3 µM) (Figure 2A). When ME was applied in the presence of TIPP[psi], the average EPSC amplitude was significantly reduced from 0.3142±0.1753 to 0.2329±0.1326 nA and IPSC amplitude was reduced from 0.6376±0.2729 to 0.3846±0.1759 nA individually (n=8, N=4, multiple paired t-test, **p<0.01, Figure 2C).

**Figure 2:**
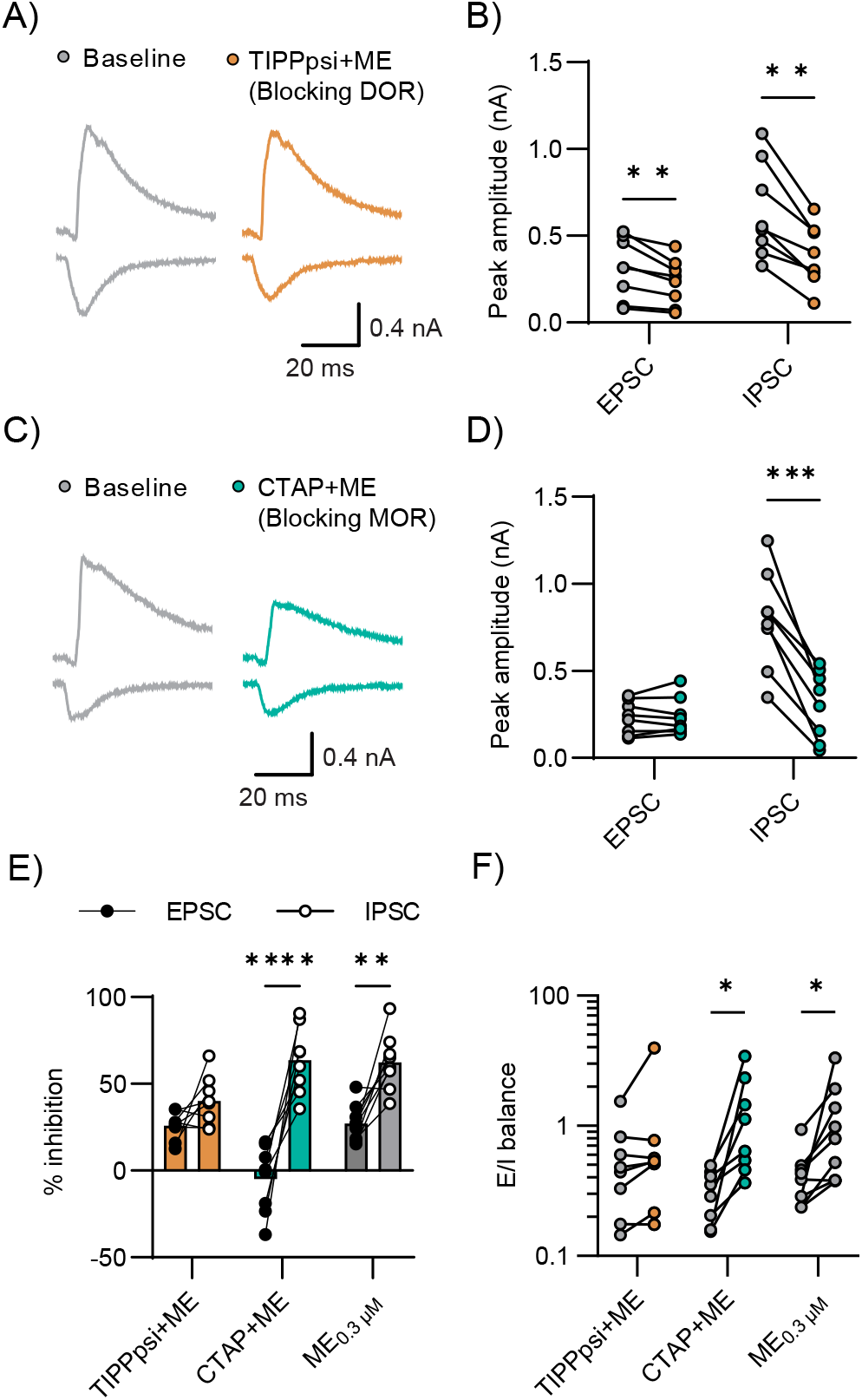
ME induced change in E/I balance was primarily mediated by DOR activation. A) Representative averaged traces of EPSCs and IPSCs recorded under baseline conditions and in the presence of 0.3 µM ME with the DOR antagonist TIPP[psi] (1 µM). B) Summary average peak EPSC and IPSC amplitudes are plotted under baseline conditions (gray) and in the presence of ME + TIPP[psi] (orange). In the presence of TIPP[psi], application of ME significantly inhibited both EPSC and IPSC (multiple paired t tests, **p<0.01). C) Representative averaged traces of EPSCs and IPSCs recorded under baseline conditions and in the presence of 0.3 µM ME with the MOR antagonist CTAP (1µM). D) Summary average peak EPSC and IPSC amplitudes are plotted under baseline conditions (gray) and in the presence of ME + CTAP (teal). In the presence of CTAP, ME significantly inhibited the IPSC but not the EPSC (multiple paired t tests, p=0.6887, ***p<0.001) E) Summary data comparing inhibition of the peak EPSC (closed) and IPSC (open) by ME + TIPP[psi] (orange), ME + CTAP (teal) and ME alone (gray) (two-way ANOVA with Šídák’s multiple comparisons test, p=0.33, ****p<0.0001,**p<0.01). F) The E/I balance in the baseline condition (TIPP[psi] or CTAP alone, respectively) versus the E/I balance in presence of ME plus TIPP[psi] or ME plus CTAP. TIPP[psi] but not CTAP blocked the effect of ME on the E/I balance (multiple Wilcoxon tests, p=0.14 and *p<0.05).

To characterize the role of DOR activation in enkephalin effects on synaptic transmission, we applied ME (0.3 µM) in the presence of the MOR antagonist CTAP (1 µM) and measured the effect on the amplitude of MThal-driven EPSCs and IPSCs in layer V pyramidal neurons in the ACC (Figure 2B). In presence of CTAP, the average amplitude of the EPSC was not reduced by enkephalin (0.2313±0.0960 nA in presence of CTAP vs 0.2377±0.1061 nA in presence of CTAP+ME, n=8, N=5, multiple paired t-test, p=0.27, Figure 2D). However, the mean amplitude of the IPSC was significantly reduced by enkephalin in presence of CTAP (0.7197±0.2848 nA in presence of CTAP vs. 0.3077±1964 nA in presence of CTAP+ME, n=8, N=5, multiple paired t-test, ***p<0.001, Figure 2D).

In the presence of TIPP[psi], there was not a statistically significant difference in the relative inhibition of EPSCs and IPSCs by ME (25.80±8.06% vs 40.24±14.33, p=0.0834, Figure 2E). In contrast, both ME alone (0.3 µM) and CTAP+ME inhibited MThal-driven IPSCs significantly more than EPSCs in the ACC: in presence of 0.3 µM ME alone, EPSCs were inhibited by 27.21±10.23%, while IPSCs were inhibited by 62.33±15.78% of control values (two-way ANOVA with multiple comparisons, p<0.0001, Figure 2E); in presence of CTAP+ME, the average inhibition of the EPSC was -4.951±19.40%, whereas the average inhibition of IPSCs was 63.53±19.26% (two-way ANOVA with multiple comparisons, ****p<0.0001). Concomitantly, The E/I balance was significantly increased in presence of 0.3 µM ME and CTAP+ME, but not in presence of TIPP[psi]+ME. ME alone increased E/I balance from 0.4178±0.2170 to 1.119±0.9849 (multiple Wilcoxon tests, *p<0.05, Figure 2F), whereas CTAP+ME increased E/I balance from 0.3134±0.1321 to 1.294±1.086 (multiple Wilcoxon tests, *p<0.05, Figure 2F). In contrast, the change in E/I balance in presence of TIPP[psi]+ME did not reach statistical significance 0.5668±0.4501 to 0.9078±1.251 (multiple Wilcoxon tests, p=0.0896, Figure 2F). These results indicate that DOR-mediated inhibition of the IPSC was required for ME to significantly shift the E/I balance towards excitation in MThal-ACC circuits.

### Selective activation of DOR but not MOR replicates the effect of enkephalin on E/I balance in thalamo-cortical synapses in the ACC

To further confirm the effects of DOR and MOR signaling to altering E/I balance in MThal-ACC synapses, we studied the effects of selective DOR-and MOR-selective agonists on EPSCs, IPSCs and E/I balance. Consistent with our previously reported results (Birdsong et al. 2019), DPDPE, a selective DOR agonist, decreased the IPSC amplitude but not the EPSC amplitude driven by optical stimulation of MThal axons in the ACC (Figure 3A). The EPSC was unchanged by DPDPE (0.3197±0.26 baseline vs. 0.3225±0.0.25 nA in DPDPE 1 µM, multiple paired t tests; p=8940), whereas the IPSC was significantly decreased (0.7688±0.35 baseline vs. 0.4032±0.27 nA in DPDPE 1 µM, multiple paired t tests; **p<0.01; n=9; N=5; Figure 3B). In contrast, DAMGO, a selective MOR agonist, inhibited both EPSCs and IPSCs (Figure 3C). In presence of 1 µM DAMGO, the average amplitude of the EPSC was reduced from 0.2814±0.26 to 0.1169±0.1 nA, and the average amplitude of the IPSC was reduced from 0.6216±0.25 to 0.1832±0.24 nA (multiple paired t tests; *p<0.05 and **p<0.01 for EPSC and IPSC respectively; n=9; N=5; Figure 3D). DPDPE inhibited IPSCs significantly more than EPSCs (wo-way RM ANOVA with multiple comparisons, the average % inhibition of EPSC and IPSC were -4.69±19.16 and 47.56±26.85, respectively; ***p<0.001, Figure 3E), whereas DAMGO inhibited both EPSCs and IPSCs to a similar extent (two-way RM ANOVA with multiple comparisons, the average % inhibition of EPSC and IPSC were 51.29±29.57 and 69.56±34.71, respectively p=0.15, Figure 3E).

**Figure 3:**
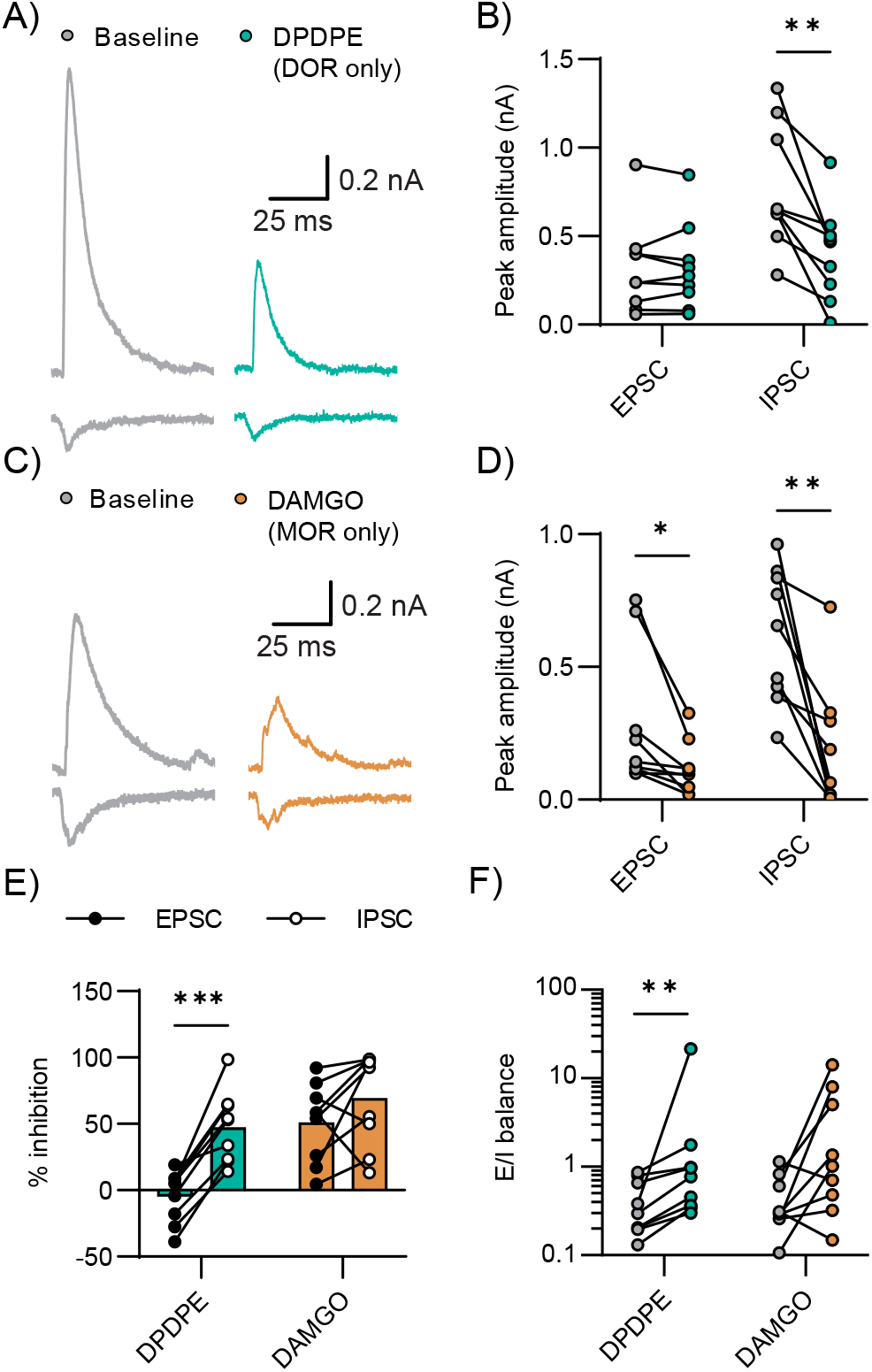
DOR agonist DPDPE mimics shift in E/I balance induced by ME. A) Example recording of EPSCs and IPSCs recorded under baseline conditions and in the presence of the MOR agonist DAMGO (1µM). B) Summary average peak EPSC and IPSC amplitudes are plotted under baseline conditions (gray) and in the presence of DAMGO (orange). DAMGO significantly inhibited both EPSC and IPSC (multiple paired t tests, *p<0.05, **p<0.01). C) Example recording of EPSCs and IPSCs recorded under baseline conditions and in the presence of the DOR antagonist DPDPE (1µM). D) Summary average peak EPSC and IPSC amplitudes are plotted under baseline conditions (gray) and in the presence of DPDPE (teal). DPDPE significantly inhibited the IPSC but did not affect the EPSC (multiple paired t tests, p=0.89 and **p<0.01). E) Summary data comparing inhibition of the peak EPSC (closed) and IPSC (open) by DAMGO (orange) and DPDPE (teal). DPDPE inhibited the EPSC and IPSC differentially, while DAMGO inhibited EPSC and IPSC proportionally (two-way ANOVA with Šídák’s multiple comparisons test, ***p<0.001 and p=0.32). F) E/I ratio in the baseline or in presence of DAMGO or DPDPE. DPDPE significantly increased E/I balance in pyramidal cells, while the effect of DAMGO on E/I balance did not reach statistical significance (multiple Wilcoxon tests, **p<0.01 and p=0.05).

Consistent with DOR activation preferentially inhibiting IPSCs, DPDPE increased the average E/I balance from 0.4153±0.28 to 3.05±6.96 (multiple Wilcoxon tests, *p<0.01, Figure 3F); while the change in E/I balance by DAMGO (0.4542±0.33 to 3.45±4.82) did not reach statistical significance (multiple Wilcoxon tests, p=0.0546, Figure 3F). These findings suggest that the impact of enkephalin on the balance between synaptic excitation and inhibition driven by MThal neurons in the ACC is primarily mediated through DOR signaling with MOR signaling perhaps further driving disinhibition.

### Enkephalin bidirectionally modulated excitatory postsynaptic potentials in ACC layer 5 pyramidal neurons

Changes in E/I balance and inhibition of excitatory transmission by ME are expected to translate into changes in postsynaptic potentials in downstream neurons; increased E/I balance is expected to increase the amplitude of postsynaptic depolarizations while decreased excitatory drive would be expected to decrease the amplitude of these depolarizations. To study the effects of ME on optically evoked excitatory postsynaptic potentials (EPSP) in ACC layer 5 pyramidal neurons, we compared two concentrations of enkephalin: 0.3 µM, a near saturating concentration, vs. 0.1 µM, a concentration that provided the maximum separation between the inhibition of the IPSC and EPSC based on our concentration-response data. Current was injected to maintain the membrane potential at -45 mV, a value that is between the EPSP and IPSP reversal potentials. QX314 was added to the internal solution to inhibit action potential firing. Baseline PSPs were evoked followed by perfusion of ME (0.1 or 0.3 µM) (Figure 4A). The near saturating concentration of 0.3 µM enkephalin produced mixed effects on the EPSP, with 7 out of 15 cells displaying a reduction in the EPSP amplitude and 4 displaying facilitation and 4 showing neither clear facilitation or reduction. On average, 0.3 µM ME did not significantly modify the amplitude of the EPSP (5.597±3.032 mV baseline, 4.483±3.377 mV in 0.3 µM ME; multiple ratios paired t tests, n=15, N=6, p=0.1503, Figure 4C). In contrast, 0.1 µM ME reliably increased the EPSP amplitude (3.087±1.626 mV baseline vs. 4.665±3.281 mV 0.1 µM ME; multiple ratios paired t-tests, n=10, N=3, *p<0.05, Figure 4C).

**Figure 4:**
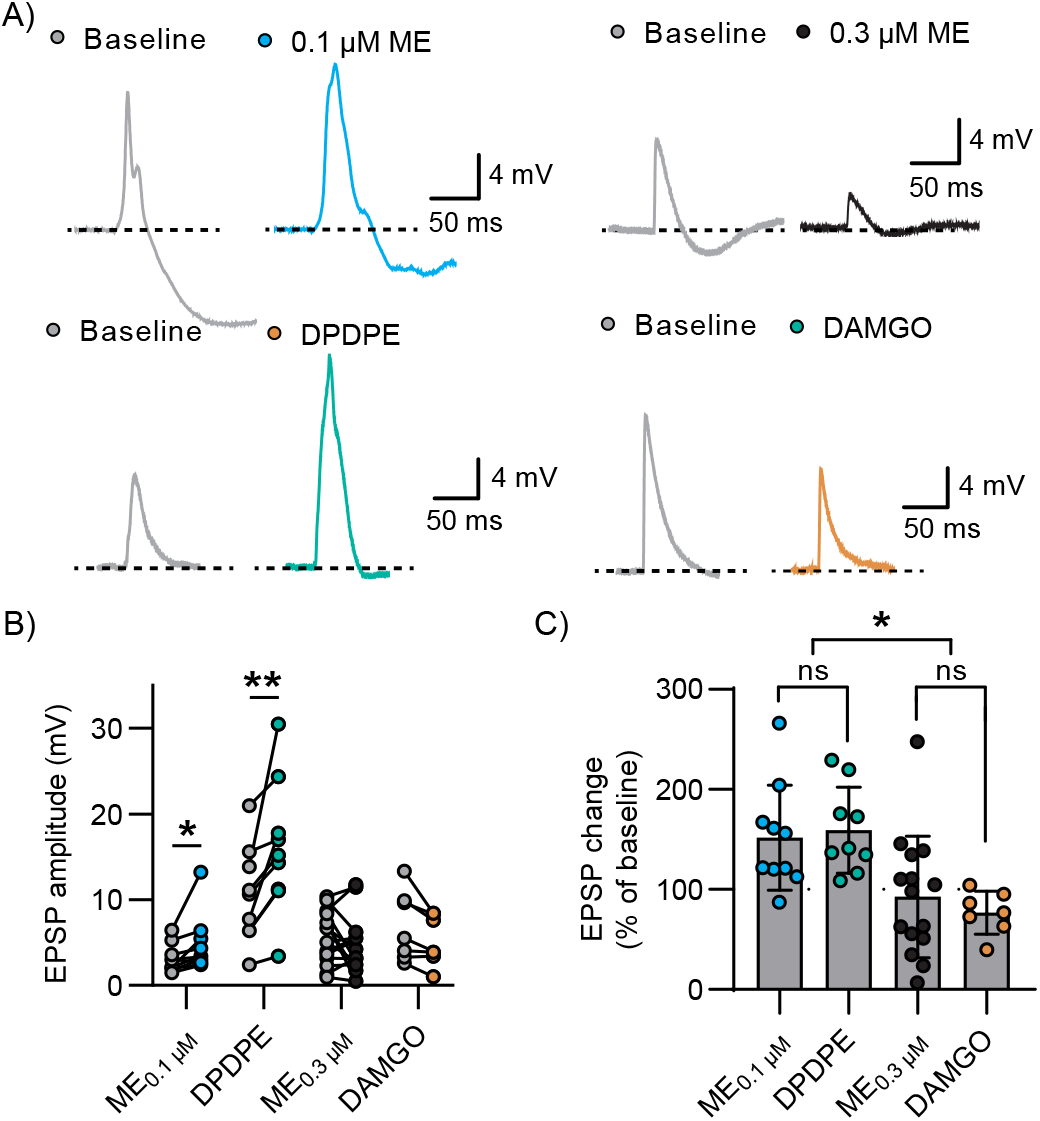
DPDPE and ME_100nM_ facilitated MThal-ACC EPSPs while DAMGO and ME_300nM_ did not. A) Representative current clamp recordings evoked by optical stimulation of MThal terminals in ACC and recorded in L5 pyramidal cells recorded under baseline conditions (gray) and in the presence of 0.1 µM ME (blue), 0.3 µM ME (black), DPDPE (teal) and DAMGO (orange). B) Summary data of EPSP peak amplitude under baseline conditions and in the presence of drug. 0.1 µM ME and DPDPE significantly increased the amplitude of the EPSP but 0.3 µM ME and DAMGO did not (multiple ratios paired t tests, *p<0.05, p<0.01, p=0.15 and p=0.09). C) Comparison of peak EPSP change plotted as the EPSP amplitude in the presence of drug as a % of baseline EPSP amplitude. 0.1 µM ME and DPDPE increased the amplitude of the EPSP in a similar fashion, while 0.3 µM ME and DAMGO did not affect the amplitude of the EPSP on average (ordinary one-way ANOVA with Tukey’s multiple comparisons, p=0.98, p=0.9, *p<0.05).

We next examined the effects of DAMGO and DPDPE on the EPSP amplitude to determine the contributions of MOR and DOR to the differential effects of enkephalin Figure 4A. DAMGO did not significantly change EPSP amplitude (6.911±4.071 mV baseline vs 5.249±2.878 mV DAMGO; ratio paired t test, n=7, N=5, p=0.0923), while DPDPE reliably increased EPSP amplitude (10.58±5.63 mV baseline vs 16.09±7.85 mV DPDPE; ratio paired t test, n=9, N=6, **p<0.01, Figure 4B).

When comparing the percent change with respect to baseline, there was no significant difference between the effect of DAMGO and ME_0.3µM_ on the EPSP amplitude (76.97±21.5% of baseline and 92.62±60.82%, respectively; ordinary one-way ANOVA with multiple comparisons, p=0.9046); whereas DPDPE and ME_0.1µM_ both increased the amplitude of the evoked EPSP to a similar extent (159.4±42.97% and 151.8±52.32% of baseline, respectively; ordinary one-way ANOVA with multiple comparisons, p=0.9875). Additionally, both DPDPE and ME_0.1µM_ increased the EPSP amplitude significantly more than either DAMGO or ME_0.3µM_ (ordinary one-way ANOVA with Tukey’s multiple comparisons test, DPDPE vs. DAMGO, p=0.0127; DPDPE vs. ME_0.3µM_, p=0.0166; ME_0.1µM_ vs. DAMGO, p=0.0231, ME_0.1µM_ vs. ME_0.3µM_, p=0.0323).

These results suggest that the relative levels of DOR and MOR expression and/or activation in thalamocortical sub-circuits dictated whether enkephalin facilitated thalamic-driven excitatory input to the ACC. By preferentially inhibiting IPSCs with either a low concentration of ME or with the DOR-selective agonist DPDPE, EPSPs were uniformly facilitated. While higher concentrations of ME still inhibited IPSCs to a greater extent than EPSC’s, decreasing excitatory drive from within the ACC eventually decreased the EPSP amplitude, nullifying this disinhibition and leading to heterogeneous effects similar to the effect of DAMGO.

## DISCUSSION

Endogenous enkephalins play important roles in pain and anxiety-related behaviors through actions on opioid receptors. The aim of this study was to understand how [Met]5-enkephalin modulates synaptic transmission between the medial thalamus and ACC. The results demonstrated that, while enkephalin inhibited both excitatory and feedforward inhibitory signaling, feedforward inhibitory signaling was preferentially suppressed. This preferential suppression of inhibitory signaling was most prominent at sub-saturating concentrations of enkephalin that led to modest inhibition of EPSCs but robust inhibition of IPSCs. The net effect of enkephalin at this modest concentration was to disinhibit ACC responses to MThal inputs, effectively increasing excitatory drive of MThal inputs onto ACC layer 5 pyramidal neurons. At higher concentrations of drug, the net effect of enkephalin was cell-dependent, with 4 of the cells showing facilitation of the EPSP and 7 showing inhibition and 4 not displaying a clear change. This biphasic effect of disinhibition and inhibition is observable at the behavioral level with many drugs including opioids. Interestingly, insulin has also been reported to have a biphasic disinhibitory and inhibitory effect on excitatory signaling, part of which appears to depend on endogenous opioids (Fetterly et al. 2021).

Endogenous enkephalin concentrations have been difficult to measure and the physiologically relevant concentrations remains unknown. Using micro dialysis and mass spectrometry, concentrations in the high picomolar range have been measured but these are likely underestimates of the true concentration due to the difficulty of collecting and isolating opioid peptides (Shen, Lada, and Kennedy 1997). Peptidase inhibitors have been shown to facilitate endogenous opioid-mediated analgesia and signaling, presumably due to increasing concentration, diffusion distance and lifetime of opioid peptides before degradation (Al-Hasani et al. 2018; Roques, Fournié-Zaluski, and Wurm 2012). These observations suggest that saturating concentrations of enkephalins are improbable under most conditions, rather, the low enkephalin concentrations that preferentially decreased inhibitory signaling, reliably increased E/I balance and disinhibited the EPSP in our experiments are likely physiologically relevant under most conditions. Additionally, the lack of any effect of perfusion of peptidase inhibitors bestatin and thiorphan (vehicle) in the concentration response data indicate that unstimulated levels of enkephalin in ACC brain slice are very low in the ACC.

Thalamic innervation is not the sole driver of ACC excitation. There are multiple glutamate inputs that may regulate ACC activity and mediate behaviors including cortico-cortical (Fillinger et al. 2017) and cortico-limbic inputs (Xu et al. 2022). Enkephalins are likely to have different effects on each input depending on the sensitivity of the afferent terminals to enkephalins, which local neuron populations are preferentially excited by these inputs and the enkephalin sensitivity of each of these interneuron populations. The net effect of enkephalins may be to bias which inputs are preferentially driving ACC activity and which inputs are suppressed. This has been previously demonstrated for modulation of prefrontal cortical (PFC) inputs and local circuits by the endogenous opioid dynorphin signaling through the kappa opioid receptor— high concentrations of dynorphin facilitated PFC responses to stimulation of ventral hippocampal inputs but suppressed responses to inputs from basolateral amygdala (Tejeda et al. 2022; Yarur et al. 2022). This biasing of the relative influence of various cortical inputs may be a common feature of opioid effects on cortical circuits that will depend on which inputs are activated, which endogenous opioids are present and their concentrations. Future studies will investigate the modulatory effects of enkephalins on signaling elicited by various inputs to the ACC.

The ACC is implicated in pain, fear, and emotional processing. Pain has been reported to alter ACC activity and endogenous opioid signaling in the ACC has been found to mediate some aspects of analgesia. This study found that modest concentrations of enkephalin may change the E/I balance of signaling within the ACC and thus either disinhibit or inhibit ACC pyramidal cells depending on concentration. Several limitations of this study can be addressed in the future to gain a clearer understanding of how opioids shape ACC function. ACC pyramidal neurons are not a homogenous population. ACC projection neurons innervate many brain regions including the periaqueductal gray, thalamus, basolateral amygdala, dorsal and ventral striatum, other cortical areas, and spinal cord and some express opioid receptors themselves.

Whether each of these ACC pyramidal cell populations receives the same relative excitatory and inhibitory innervation. However, synaptic inputs to different classes of cortical pyramidal neurons may be differentially altered in response to challenges such as chronic pain, suggesting that there is likely heterogeneity within ACC subcircuits (Meda et al. 2019). This heterogeneity may explain some of the variability in EPSP responses seen in our results.

Further heterogeneity may exist due to differential opioid receptor expression in subsets of pyramidal neurons themselves. Some ACC pyramidal neurons express opioid receptors and their activation has been reported to hyperpolarize these cells, introducing further heterogeneity in the output of pyramidal neurons in response to synaptic inputs and enkephalins (Tanaka and North 1994).

However, in our recording configuration, hyperpolarization or inward currents would not be clearly observed due to the internal solutions used and current offsets used. Overall, the present study provides a framework for studying and understanding how endogenous opioids can act within the ACC to bias responses of ACC subcircuits to cortical inputs. It also demonstrates that receptor expression levels, endogenous opioid concentration and relative selectivity or lack thereof of opioid/receptor interactions can affect circuit output in complex but predictable ways.

## AUTHOR CONTRIBUTIONS

ERAH: Conceptualization, Formal Analysis, Investigation, Visualization, Methodology, Writing

WTB: Conceptualization, Formal Analysis, Funding Acquisition, Resources, Supervision, Visualization, Methodology, Writing

## ACKNOWLEDGEMENTS

This work was supported by R01DA042779 (WTB)

